# Long-term antitumor CD8^+^ T cell immunity induced by endogenously engineered extracellular vesicles

**DOI:** 10.1101/2021.02.05.429897

**Authors:** Flavia Ferrantelli, Francesco Manfredi, Chiara Chiozzini, Eleonora Olivetta, Andrea Giovannelli, Patrizia Leone, Maurizio Federico

## Abstract

We developed a novel approach to induce antigen-specific CD8^+^ T cytotoxic lymphocyte (CTL) immunity based on *in vivo* engineering of extracellular vesicles (EVs). This is an innovative vaccination approach employing a DNA vector expressing a mutated HIV-1 Nef protein (Nef^mut^) that has lost the anti-cellular effects typical of the wild-type isoform, meanwhile showing an unusual efficiency of incorporation into EVs. This function persists even when foreign antigens are fused to its C-terminus. In this way, Nef^mut^ traffics large amounts of antigens fused to it into EVs spontaneously released by cells expressing the Nef^mut_^based DNA vector. We previously provided evidence that the inoculation in mice of a DNA vector expressing the Nef^mut^/HPV16-E7 fusion protein induced an E7-specific CTL immune response as detected 2 weeks after the second immunization. In an effort to optimize the anti-HPV16 CD8^+^ T cell immune response, we found that the co-injection of DNA vectors expressing Nef^mut^ fused with E6 and E7 generated a stronger anti-HPV16 immune response compared to that we observed in mice injected with the single vectors. When TC-1 cells, i.e., a tumor cell line co-expressing E6 and E7, were implanted before immunization, all mice survived until day 44, whereas no mice injected with either void or Nef^mut_^expressing vectors survived until day 32 after tumor implantation. A substantial part of mice (7 out of 12) cleared the tumor. When cured mice were re-challenged with a second sub cute implantation of TC-1 cells, and followed for additional 135 days, whereas none of them developed tumors. Both E6- and E7-specific CD8^+^ T immunity was still detectable at the end of the observation time.

Hence, the immunity elicited by engineered EVs, besides curing already developed tumors, is strong enough to guarantee the resistance to additional tumor attack. This results is of relevance for therapy against both metastatic and relapsing tumors.

## Introduction

Eukaryotic cells spontaneously release vesicles of different sizes. Extracellular vesicles (EVs) are classified as apoptotic bodies (1-5 μm), microvesicles (50-1,000 nm), and exosomes (50-200 nm) (1). Microvesicles (also referred to as ectosomes) shed by plasma membrane, whereas exosomes are released after inward invagination of endosome membranes and formation of intraluminal vesicles. Healthy cells constitutively release both exosomes and microvesicles, together referred to as EVs hereinafter. EVs are an important mean of intercellular communication by transporting their cargo, such as DNAs, RNAs, proteins, and lipids, from the producer cell to the recipient one (2, 3).

EVs are abundant, stable, and highly bioavailable to tissues *in vivo*. They find potential applications as diagnostic biomarkers, therapeutics, drug delivery vehicles, and functional cosmetics. Several EV-based anticancer immunotherapies have been under clinical trials for different indications (4, 5). Despite high expectations, clinical trials have not yet confirmed therapeutic application of *in vitro* engineered EVs, mostly because a number of drawbacks associated with functional reproducibility and loading of specific cargoes (6).

Two types of cancer immunotherapy have emerged so far as being the most promising: i) T cell-based cancer immunotherapy, including active vaccination and adoptive cell transfer, and ii) immune modulation through monoclonal antibodies referred to as immune checkpoint blockers (ICBs) (7).

Cancer vaccines rely on induction of antitumor immunity using whole or part of tumor antigens. In this manner, a *de novo* antitumor immunity can be established, and both potency and breadth of pre-existing immunity can be widened. Different approaches include the use of synthetic peptides from tumor antigens (allogenic and autologous), dendritic cell (DC)-based vaccines, and genetic vaccines (DNA/RNA/virus/bacterial) (8).

Cancer cells express a burden of new antigens as a result of their intrinsic genetic instability typical of malignant transformation and/or of the expression of the etiologic cancer agents, as in the case of virus-induced malignancies. In non-virus-induced cancers, transformed cells can produce antigens to which the host is basically tolerant (tumor-associated self-antigens. In addition, cancer cells can express antigens to which the host does not develop tolerance, being however the immune response not effective enough to counteract the cancer cell growth. They include the so-called “tumor specific neo-antigens” as well as antigens normally produced in immune-privileged tissues, e.g., cancer-testis antigens. Hence, establishing a method to induce an adaptive immune response against both tolerogenic and non-tolerogenic TAAs would be of great relevance for the design of novel antitumor therapeutic approaches.

An explosive growth of interest in cancer immunotherapy occurred in the last decade mainly due to approval of new clinical protocols based on the use of monoclonal antibodies, referred to as ICBs (9, 10), fostering the pre-existing immune response against both tumor-associated antigens (TAAs) and neo-antigens. The success of immunotherapies based on ICBs definitely proved that cancer can be treated and cured by manipulating the immune system. However, this strategy still suffers from some limitations, including intrinsic and/or acquired resistance, and development of hyper-immune activation, which can associate with immune-related adverse events affecting several organs including skin, gut, heart, lungs, and bone (11).

DNA vaccination has many potential advantages (12). DNA molecules (usually plasmids) express the antigen of interest under the control of a strong promoter are transferred to cells of a vaccine recipient. Intracellularly produced antigens are presented to the recipient’s immune system, resulting in both humoral and cellular immune responses that may protect against disease in preclinical models of cancer, infectious diseases, and autoimmunity. Nevertheless, to date the efficacy of DNA vaccines in clinical trials has been disappointing, and it is uncertain whether the high expectations associated with DNA vaccines will be fulfilled. Anyway, preclinical and clinical studies have yielded many safety data (13).

We developed a vaccine platform based on the high levels of uploading into extracellular vesicles (EVs) of a Human Immunodeficiency virus-1 Nef mutant, referred to as Nef^mut^ (14). In the Nef^mut_^based biotechnology platform, the antigen of interest is expressed by a DNA vector as product of fusion to Nef^mut^. Upon intramuscular injection, the antigen is incorporated into the EVs which are spontaneously released by muscle cells. These EVs freely circulate into the body reaching also compartments distal from the injection site. When they enter professional APCs, the Nef^mut_^antigen product of fusion is cross-presented, thereby inducing antigen-specific CTLs (15–17). On the contrary, no humoral response is produced most likely as a consequence of the incorporation of Nef^mut_^based fusion products into EVs, where they remain hidden and protected from external environment.

Here, data regarding both duration and efficacy against tumor challenge and re-challenge of an anti-HPV16 vaccine based on Nef^mut^ *in vivo*-engineered EVs are reported.

## Materials and Methods

### DNA vector synthesis

The pTargeT (Invitrogen, Thermo Fisher Scientific) vectors expressing Nef^mut^, Nef^mut^/HPV16-E6 and Nef^mut^/HPV16-E7 were already described (15, 18). Both E6 and E7 ORFs were synthesized by Eurofins Genomics Germany GmbH. Kozak sequences were inserted at 5’ end, and ORFs were inserted in the *Not* I and *Apa* I sites of the pTargeT vector polylinker. A 6×His tag sequence (i.e., 5’ CACCATCACCATCACCAT 3’) was included at the 3’ end just before the stop codon.

### Cell cultures and transfection

Human embryonic kidney (HEK)293T cells (ATCC, CRL-11268) were grown in DMEM (Gibco) plus 10% heat-inactivated fetal calf serum (FCS, Gibco). Transfection assays were performed using Lipofectamine 2000 (Invitrogen, Thermo Fisher Scientific).

### EV isolation

Cells transfected with vectors expressing the Nef^mut_^based fusion proteins were washed 24 h later, and reseeded in medium supplemented with EV-deprived FCS. The supernatants were harvested from 48 to 72 h after transfection. EVs were recovered through differential centrifugations (19) by centrifuging supernatants at 500×*g* for 10 min, and then at 10,000×*g* for 30 min. Supernatants were harvested, filtered with 0.22 μm pore size filters, and ultracentrifuged at 70,000×*g* for l h. Pelleted vesicles were resuspended in 1×PBS, and ultracentrifuged again at 70,000×*g* for 1 h. Finally, pellets containing EVs were resuspended in 1:100 of the initial volume.

### Western blot analysis

Western blot analyses of both cell lysates and EVs were carried out after resolving samples in 10% sodium dodecyl sulfate-polyacrylamide gel electrophoresis (SDS-PAGE). In brief, the analysis on cell lysates was performed by washing cells twice with 1×PBS (pH 7.4) and lysing them with 1x SDS-PAGE sample buffer. Samples were resolved by SDS-PAGE and transferred by electroblotting on a 0.45 μM pore size nitrocellulose membrane (Amersham) overnight using a Bio-Rad Trans-Blot. For western blot analysis of EVs, they were lysed and analyzed as described for cell lysates. For immunoassays, membranes were blocked with 5% non-fat dry milk in PBS containing 0.1% Triton X-100 for 1 h at room temperature, then incubated overnight at 4 °C with specific antibodies diluted in PBS containing 0.1% Triton X-100. Filters were revealed using 1:1,000-diluted sheep anti-Nef antiserum ARP 444 (MHRC, London, UK), 1:500-diluted anti-β-actin AC-74 mAb from Sigma, and 1:500 diluted anti-Alix H-270 polyclonal Abs from Santa Cruz.

### Mice immunization

Six-weeks old C57 Bl/6 female mice were obtained from Charles River and placed in the Central Animal Facility of the ISS, as approved by the Italian Ministry of Health, authorization n. 950/2018. The DNA vector preparations were diluted in sterile 0.9% saline solution. Both quality and quantity of the DNA preparations were checked by 260/280 nm absorbance and electrophoresis assays. Mice were anesthetized with isoflurane as prescribed in the Ministry authorization. Each inoculum volume was injected into both quadriceps. Immediately after inoculation, electroporation was applied at the site of injection through the Agilepulse BTX device using a 4-needle array 4 mm gap, 5 mm needle length, BD, with the following parameters: 1 pulse of 450 V for 50 μsec; 0.2 msec interval; 1 pulse of 450 V for 50 μsec; 50 msec interval; 8 pulses of 110 V for 10 msec with 20 msec intervals. Immunizations were repeated identically 14 days later. For immunogenicity studies, fourteen days after the last immunization mice were sacrificed by cervical dislocation as recommended by the Ministry authorization protocol. Spleens were then explanted and placed into a 2 mL Eppendorf tubes filled with 1 mL of RPMI 1640 (Gibco), 50 μM 2-mercaptoethanol (Sigma). PBMCs were recovered from blood samples obtained by retro orbital bleeding. Red blood cells were eliminated through incubation with ACK lysing buffer (Gibco).

### IFN-γEliSpot analysis

For the IFN-γ EliSpot assay, 2.5×10^5^ live cells were seeded in each microwell. Cultures were run in triplicate in EliSpot multiwell plates (Millipore, cat n. MSPS4510) pre-coated with the AN18 mAb against mouse IFN-γ (Mabtech) in RPMI 1640 (Gibco) plus 10% FBS (Gibco) for 16 h in the presence or not of 5 μg/mL of the following peptides (Table 1): E6 18-26: KLPQLCTEL (20); 50-57: YDFAFRDL (20); 109-117: RCINCQKPL (21); 127-135: DKKQRFMNI (20), and E7: 49-57: RAHYNIVTF (20); 67-75: LCVQSTHVD (21). As a negative control, 5 μg/mL of the H2-K^b^-binding HCV-NS3 specific peptide ITQMYTNV (22) were used. More than 70% pure preparations of the peptides (recommended peptide purity for performing an EliSpot assay) were obtained from either UFPeptides, Ferrara, Italy, or JPT, Berlin, Germany. For cell activation control, cultures were treated with 10 ng/mL PMA (Sigma) plus 500 ng/mL of ionomycin (Sigma). After 16 hours, cultures were removed, and the wells incubated with 100 μL of 1 μg/ml of the R4-6A2 biotinylated anti-IFN-γ (Mabtech) for 2 hours at r.t. Wells were then washed and treated for 1 hour at r.t. with 1:1000 diluted streptavidine-ALP preparations from Mabtech. After washing, spots were developed by adding 100 μL/well of Sigma Fast BCIP/NBT. The spot-forming cells were finally analyzed and counted using an AELVIS EliSpot reader.

### Intracellular cytokine staining (ICS)

Splenocytes were seeded at 2×10^6^/mL in RPMI medium, 10% FCS, 50 μM 2-mercaptoethanol (Sigma), and 1 μg/mL brefeldin A (BD Biosciences). Control conditions were carried out either by adding 10 ng/ml PMA (Sigma) and 1 μg/mL ionomycin (Sigma), or with unrelated peptides. After 16 hours, cultures were stained with 1 μl of LIVE/DEAD Fixable Aqua Dead Cell reagent (Invitrogen ThermoFisher) in 1 mL of PBS for 30 minutes at 4 °C and washed twice with 500 μL of PBS. To minimize nonspecific staining, cells were pre-incubated with 0.5 μg of Fc blocking mAbs (i.e., anti-CD16/CD32 antibodies, Invitrogen/eBioscience) in 100 μL of PBS with 2% FCS for 15 minutes at 4 °C. For the detection of cell surface markers, cells were stained with 2 μL of the following Abs: FITC conjugated anti-mouse CD3, APC-Cy7 conjugated anti-mouse CD8a, and PerCP conjugated anti-mouse CD4 (BD Biosciences) and incubated for 1 hour at 4 °C. After washing, cells were permeabilized and fixed through the Cytofix/Cytoperm kit (BD Biosciences) as per the manufacturer’s recommendations, and stained for 1 hour at 4 °C with 2 μl of the following Abs: PE-Cy7 conjugated anti-mouse IFN-γ (BD Biosciences), PE conjugated anti-mouse IL-2 (Invitrogen eBioscience), and BV421 anti-mouse TNF-α (BD Biosciences) in a total of 100 μL of 1× Perm/Wash Buffer (BD Biosciences). After two washes, cells were fixed in 200 μL of 1× PBS/formaldehyde (2% v/v). Samples were then assessed by a Gallios flow cytometer and analyzed using Kaluza software (Beckman Coulter).

Gating strategy was as follows: live cells as detected by Aqua LIVE/DEAD Dye vs. FSC-A, singlet cells from FSC-A vs. FSC-H (singlet 1) and SSC-A vs SSC-W (singlet 2), CD3 positive cells from CD3 (FITC) vs. SSC-A, CD8 or CD4 positive cells from CD8 (APC-Cy7) vs. CD4 (PerCP). The CD8^+^ cell population was gated against APC-Cy7, PE, and BV421 to observe changes in IFN-γ, IL-2, and TNF-α production, respectively. Boolean gates were created in order to determine any cytokine co-expression pattern.

### Tumor challenge and re-challenge

Six-weeks old C57 Bl/6 female mice were obtained from Charles River and placed in the Central Animal Facility of the ISS. TC-1 cells (a kind gift of Prof. Wu, Johns Hopkins University, Boston, MA) were prepared at 2×10^6^ cells/mL in 1×PBS, and s.c. inoculated into mice with a volume of 100 μL. At the time of immunization, mice were anesthetized with isoflurane as prescribed in the Ministry authorization. Immunizations were carried out as here above described and repeated after 10 days. To collect peripheral blood mononuclear cells (PBMCs) for evaluating the immune responses, seven days after the second immunization 200 μL of blood were collected from each mouse through retro orbital bleeding. Tumor growth was monitored daily by visual inspection, palpation, and measurement of diameters by an electronic caliper, and volumes calculated as (length×width^2^)/2. Mice were sacrificed by cervical dislocation as recommended by the Ministry authorization protocol if in poor health or as soon as tumors reached the size of 1 cm^3^.

Nineteen weeks after the last immunization, six mice that had recovered from tumor growth were rechallenged with the same cells under the same experimental modalities used for the first implantation. Briefly, TC-1 cells were prepared at 2×10^6^ cells/mL in 1×PBS, and s.c. inoculated into mice (100 μL). Hence, a total of 2×10^5^ TC-1 cells were implanted s.c. in the opposite side respect to the previous cell implantation. As a control, two age-matched (i.e., seven months old) naïve mice were injected with TC-1 cells with identical modalities.

Tumor growth was monitored as above described. Mice were sacrificed by cervical dislocation as soon as the tumor reached the size of 1 cm^3^, or at the end of experiment, i.e., 19 weeks after tumor cell reimplantations.

### Statistical analysis

When appropriate, data are presented as mean + standard deviation (SD).

## Results

### Additive anti-HPV16 immunogenicity in mice injected with DNA vectors separately expressing E6 and E7 fused with Nef^mut^

Very recently, we provided evidence that the co-injection of up to three distinct DNA vectors expressing SARS-CoV-2 antigens fused with Nef^mut^ generated an additive CD8^+^ T cell immune response, in the absence of evident negative interferences among the different immunogens (23). We aimed at assessing whether a similar additive effect would take place with HPV16 antigens. To this aim, two DNA vectors expressing Nef^mut^/E6 and Nef^mut^/E7 were considered. In these DNA vectors the HPV16-related ORFs were optimized for the expression in eukaryotic cells, and the domains involved in the pathogenic interaction with cell protein targets were inactivated (24, 25).

Both vectors were proven to express fusion proteins which are efficiently uploaded into EVs (Fig. 1A). DNA vectors were i.m. injected in mice either singularly or in combination. Fourteen days after the second immunization, the CD8^+^ T cell immune response was evaluated in terms of antigen-specific activation through both IFN-γ EliSpot assay and ICS analysis.

**Figure 1.**
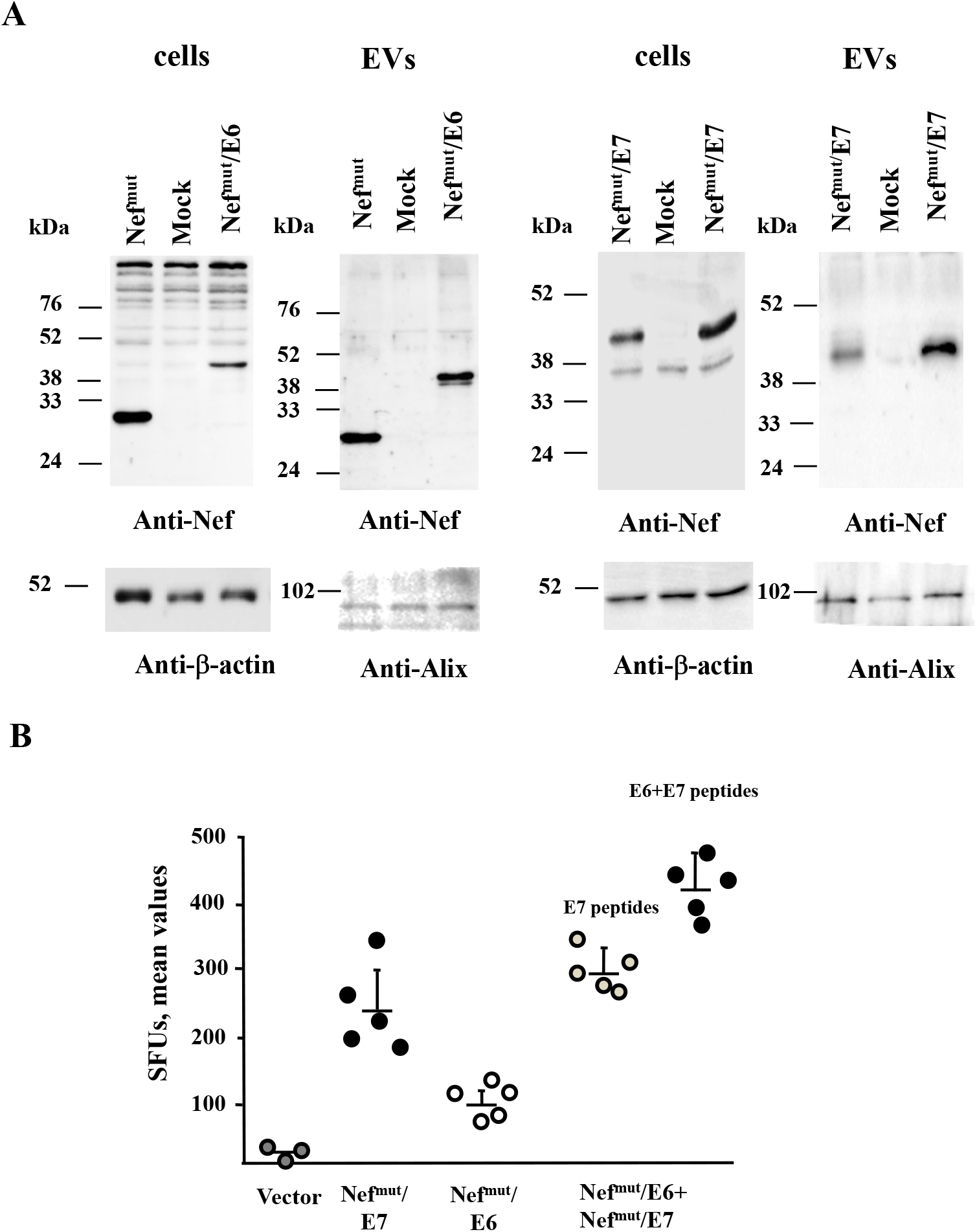
HPV16-E6 and -E7-specific CD8^+^ cell immunity induced in mice co-injected with Nef^mut^/E6 and Nef^mut^/E7 DNA vectors. A. Detection of Nef^mut_^related fusion proteins in transfected cells and EVs. Western blot analysis from 30 μg of cell lysates from 293T cells transfected with DNA vectors expressing Nef^mut^/E6 and Nef^mut^/E7, and equal volumes of buffer where purified EVs were resuspended after differential centrifugations of the respective supernatants. As control, conditions from mock-transfected cells as well as cells transfected either with Nef^mut^ or Nef^mut^/E7 were included. Polyclonal anti-Nef Abs served to detect Nef^mut_^based products, while β-actin and Alix were markers for cell lysates and exosomes, respectively. Nef protein products are indicated by arrows. Molecular markers are given in kDa. B. CD8^+^ T cell immune response induced in C57 Bl/6 mice inoculated with the DNA vectors expressing Nef^mut^/E6 and Nef^mut^/E7 either singularly or in combination. As control, mice were inoculated with the void vector. At the time of sacrifice, 2.5×10^5^ splenocytes were incubated o.n. with or without 5 μg/ml of either unrelated, E6, E7, and E6+E7-specific peptides in triplicate IFN-γ EliSpot microwells. For each mouse, shown are the numbers of IFN-γ spot-forming units (SFU) as mean values of triplicates after subtraction of values from wells treated with unrelated peptides. Intragroup mean values+SD are also reported. The E7-specific immune response was evaluated also in co-injected mice.

We observed general stronger CD8^+^ T cell responses against E7 compared to E6. When the two vectors were co-injected, additive immune responses were generated (Fig. 1B), thus excluding possible functional interference. This conclusion was also supported by the evidence that in co-injected animals not only was the E7-specific CD8^+^ T cell response not reduced, but resulted slightly increased compared to that of mice injected with the Nef^mut^/E7-expressing vector alone (Fig. 1B).

The induction of antigen-specific polyfunctional lymphocytes is considered hallmark of efficacy for CD8^+^ T cell immune response. We evaluated the levels of E6- and/or E7-specific CD8^+^ T cells expressing IFN-γ, IL-2, and TNF-α by ICS after overnight cultivation of splenocytes with specific nonamers. Consistently with data obtained from IFN-γ EliSpot assays, the immunization with the Nef^mut^/E7-expressing vector resulted in higher percentages of IFN-γ producing CD8^+^ T cells compared to those from Nef^mut^/E6 immunized mice (Fig. 2A). The highest percentages of IFN-γ producing CD8^+^ T cells was observed with splenocytes from mice injected with the two DNA vectors (Fig. 2A). Similar results were obtained with the analysis of IL-2 and TNFα. Most important, the percentages of triple-positive (i.e., polyfunctional) antigen-specific CD8^+^ T cells increased in co-injected mice compared to mice injected with single vectors (Fig. 2B).

**Figure 2.**
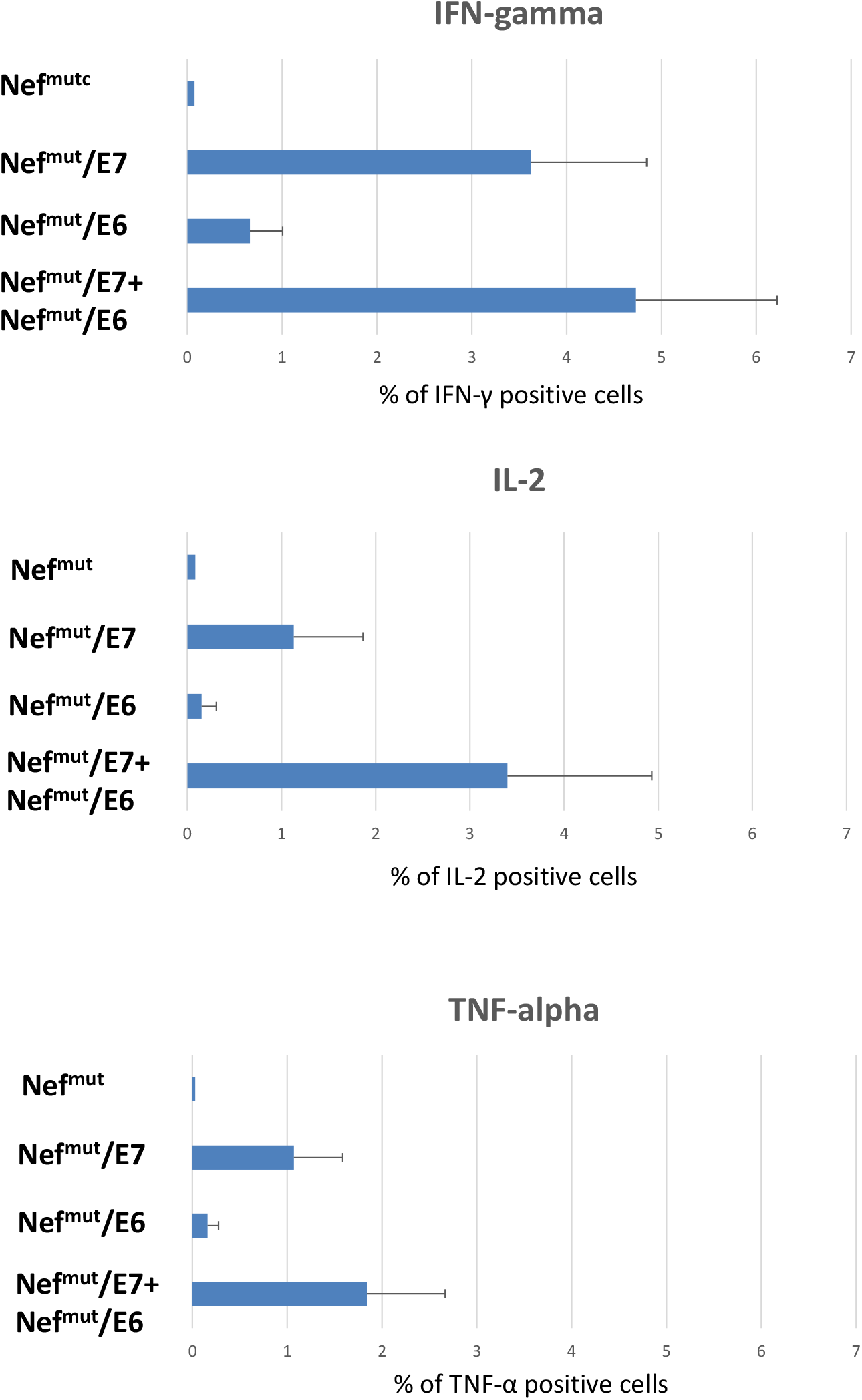
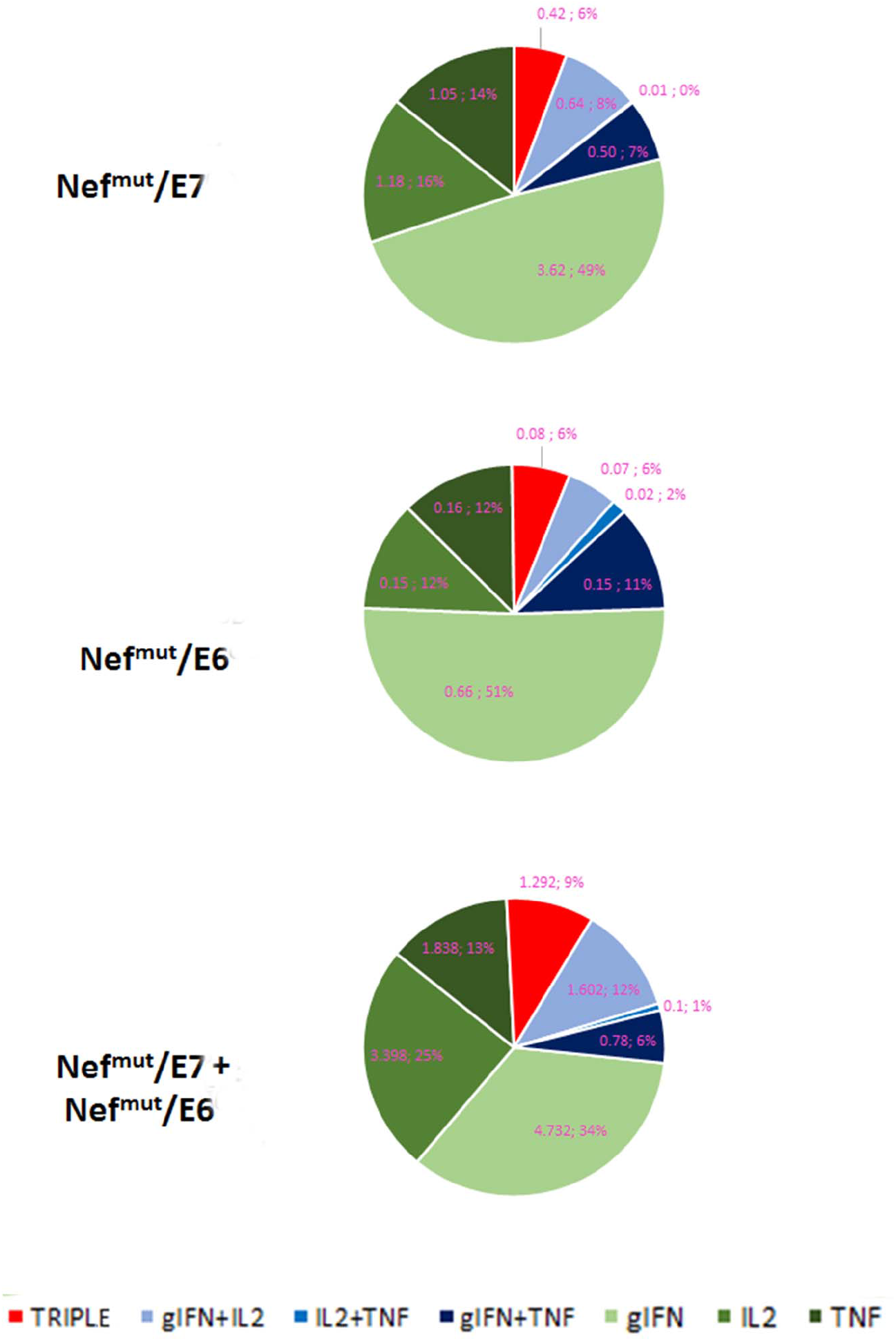
Intracellular cytokine staining analysis on splenocytes from injected mice. A. Percentages of IFN-γ, IL-2 and TNF-α accumulating CD8^+^ T cells within total CD8^+^ T cells from cultures of splenocytes isolated from each mouse injected with the indicated DNA vectors. Shown are mean values +SD of the absolute percentages of positive CD8^+^ T cells within total CD8^+^ T cells from cultures treated with specific peptides subtracted values detected in CD8^+^ T cells from cultures treated with unrelated peptide. B. Intragroup mean percentages of IFN-γ, IL-2 and TNF-α positive CD8^+^ T cells within total CD8^+^ T cells from cultures of splenocytes isolated from each mouse injected with the indicated DNA vectors, and of the CD8^+^ T cell sub-populations accumulating all possible combinations of cytokines. Shown are both absolute and relative mean percentages of positive CD8^+^ T cells within total CD8^+^ T cells from cultures treated with specific peptides subtracted values detected in CD8^+^ T cells from cultures treated with unrelated peptide. Values detected with splenocytes from mice injected with control vector were below the sensitivity threshold of the assay.

We concluded that an optimal CD8^+^ T cell immune response can be achieved using the combined immunization with Nef^mut^/E6- and Nef^mut^/E7-expressing DNA vectors.

### HPV16-E6 and –E7-specfic CD8^+^ T cell immune response in mice immunized after tumor implantation

Next, the CD8^+^ T cell immune response against both E6 and E7 was tested in mice immunized after the s.c. implantation of TC-1 tumor cells. The actual expression of both HPV16 E6 and -E7 genes in TC-1 cells was confirmed by qRT-PCR assay (not shown). C57 Bl/6 mice were implanted with 2×10^5^ TC-1 cells and, 10 days thereafter, the first immunization was carried out in mice bearing palpable tumors. Mice were injected with both Nef^mut^/E6- and Nef^mut^/E7-expressing vectors or, as control,: i) void DNA vector; ii) a vector expressing Nef^mut^ alone, and iii) a combination of vectors expressing E6 and E7. As expected, neither E6 nor E7 associated with EVs, as shown by the western blot comparative analysis including a His-tagged single-chain antibody (scFv) fused with Nef^mut^ (26) (Fig. 3A). The immunizations were repeated after a week, and after additional seven days, 200 μL of blood were recovered to evaluate the immune responses.

**Figure 3.**
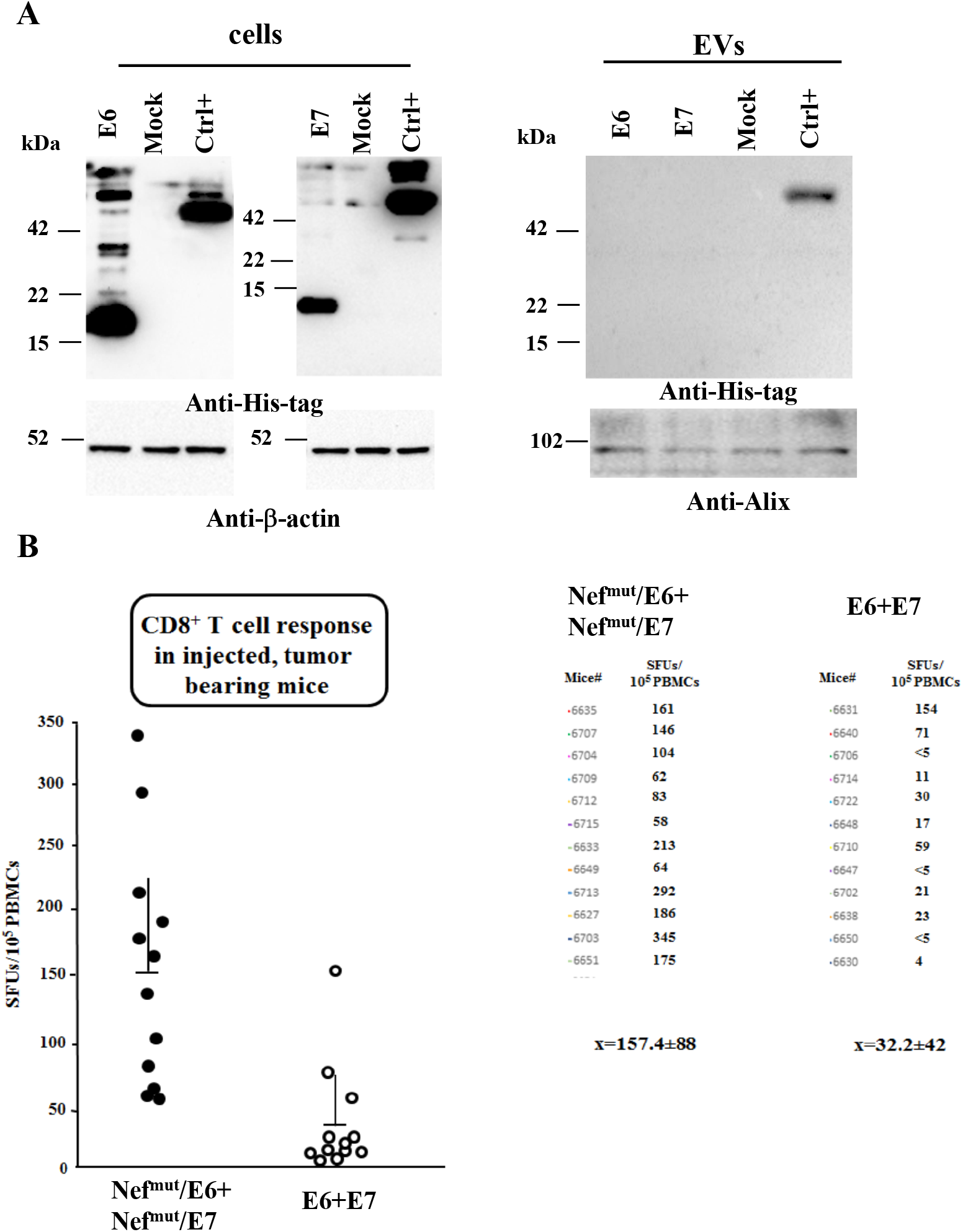
HPV16 E6- and E7-specific CD8^+^ T cell immunity induced in tumor-bearing co-injected mice. A. Analysis of both HPV16 E6 and E7 products in transfected cells and respective EVs. Western blot analysis from 30 μg of cell lysates from 293T cells transfected with DNA vectors expressing the indicated HPV16 ORFs (left panels), and equal volumes of buffer where purified EVs were resuspended after differential centrifugations of the respective supernatants (right panels). As control, conditions from mock-transfected cells as well as cells transfected with Nef^mut^ fused with a scFv (Nef^mut^ GO) including an His-tag at its C-terminus were included. Polyclonal anti-His-tag Abs served to detect both HPV16-related and Nef^mut-^ GO products, while β-actin and Alix were revealed as markers for cell lysates and EVs, respectively. Relevant protein products are highlighted. Molecular markers are given in kDa. B. CD8^+^ T cell immune response in C57 Bl/6 mice injected s.c. with 2×10^5^ TC-1 cells, and then inoculated+EP with 10 μg of both DNA vectors expressing E6 and E7 either alone or fused with Nef^mut^. PBMCs were isolated after retro orbital bleeding, and then incubated o.n. with or without 5 μg/ml of either unrelated (not shown), E6 and E7-specific nonamers in triplicate IFN-γ EliSpot microwells. Shown are the number of IFN-γ spot-forming units (SFU)/10^5^ PBMCs as mean values of triplicates. On the right, the SFU values are associated with each mouse. Intragroup mean values ± SD are also reported.

The analysis of E6- and E7-specific CD8^+^ T cell immune responses carried out by IFN-γ Elispot assay on PBMCs shown an immune response about 5-fold more potent in mice injected with DNA vectors expressing the HPV16 proteins fused with Nef^mut^ than detected in mice injected with E6 and E7-expressing vectors without Nef^mut^ (Fig. 3B). In this experimental setting, the sole implantation of the E6- and E7-expressing TC-1 cells did not result in a CD8^+^ T cell immune response, as indicated by the analysis carried out with PBMCs from mice injected with either void or Nef^mut_^expressing DNA vectors (not shown).

Overall, immune responses to Nef^mut_^based products were in line with previous observations in tumor-free mice.

### Therapeutic antitumor effect in mice co-injected with vectors expressing both E6- and E7-based fusion proteins

The tumor size of injected mice was evaluated over the time. Tumors implanted in mice injected with void or Nef^mut-^expressing vectors grew in a very quick and uncontrolled way, rapidly leading the mice to death (Fig. 4A). Tumor growth was less rapid in mice injected with vectors expressing E6 and E7. In this group, the mouse showing the strongest CD8^+^ T cell response remained tumor-free over the time, while in the remainders the tumor led the mice to reach the maximum tumor size allowed before euthanasia (i.e., 1 cm^3^) within 60 days (Fig. 4A). Conversely, seven of the twelve mice immunized with Nef^mut^/E6 plus Nef^mut^/E7-expressing vectors were cured and remained tumor-free over the time. In particular, at day 35 after tumor implantation (i.e., when all mice of control groups had been euthanized) no or very limited tumor growth was observed in mice co-injected with Nef^mut^/E6 plus Nef^mut^/E7 DNA vectors. At day 65 after tumor implantation, when only 1 mouse of 12 mice injected with DNA vectors expressing E6 and E7 was alive, all mice immunized with Nef^mut^/E6 plus Nef^mut^/E7 still survived. As shown by the survival curve (Fig. 4B), seven mice of the group immunized with DNA vectors expressing Nef^mut^/E7 and Nef^mut^/E6 remained tumor-free throughout.

**Figure 4.**
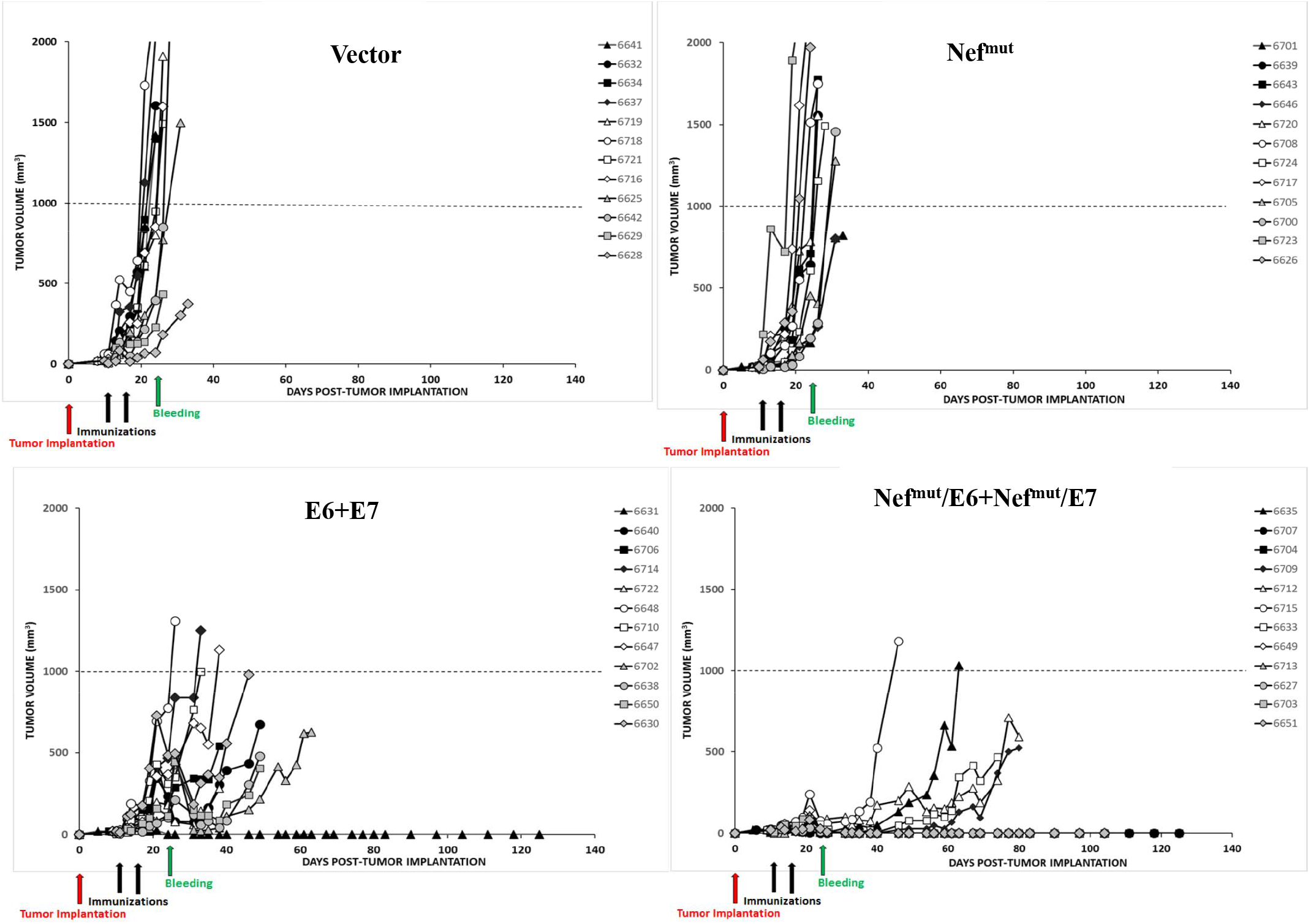
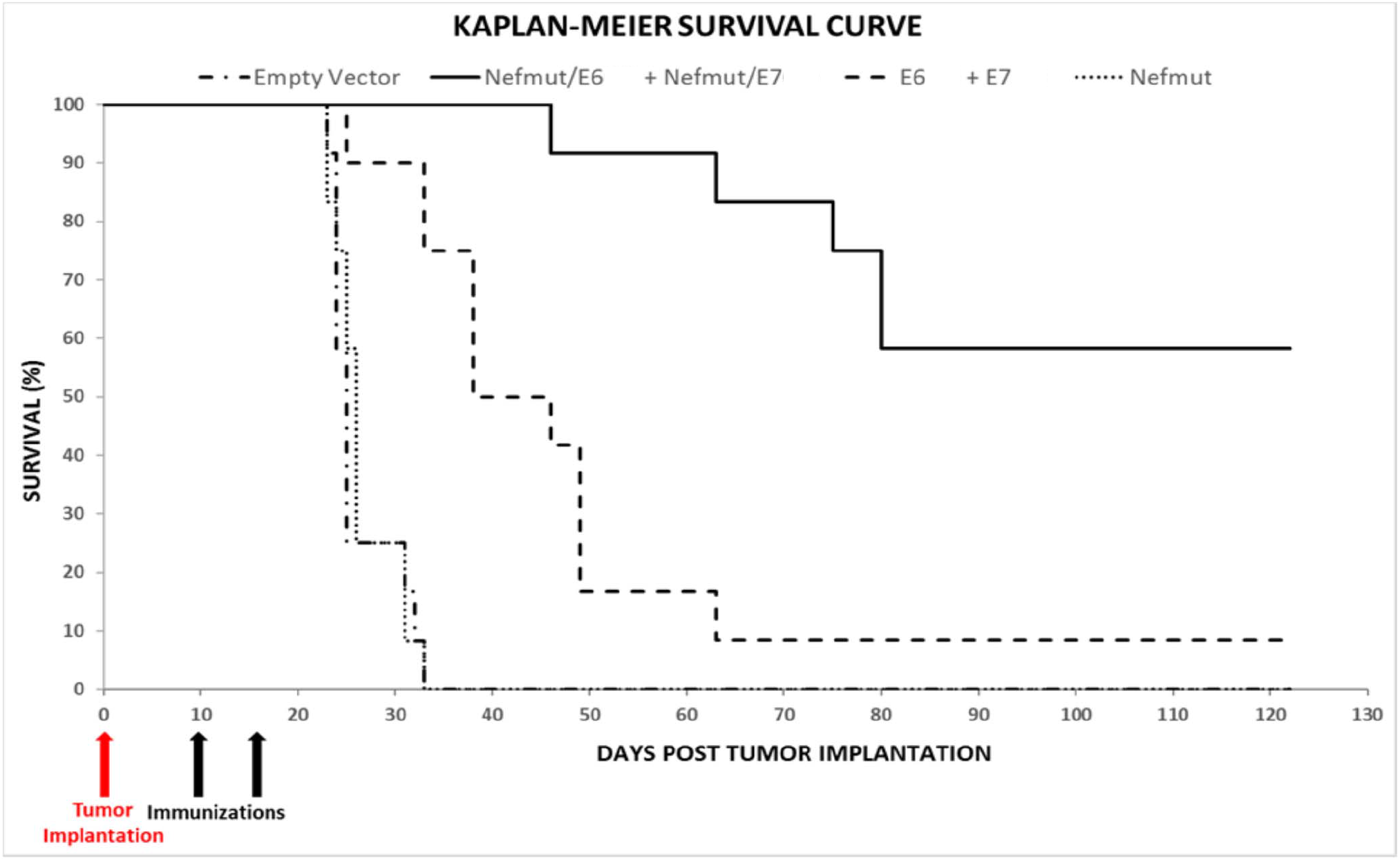
Antitumor therapeutic effect induced by i.m. injection of both Nef^mut^/E6 and Nef^mut^/E7 DNA vectors. A. Tumor growth curves. C57 Bl/6 mice (12 per group) were challenged with 2×10^5^ TC-1 cells and, the day after the tumor appearance, i.e., when tumor masses became detectable by palpation, co-inoculated with DNA vectors expressing Nef^mut^/E6, Nef^mut^/E7, E6, and E7, or, as control, with either Nef^mut^ or empty vector. The DNA inoculations were repeated at day 17 after tumor cell implantation, and the growth of tumor mass was followed over the time. Shown are the data referred to each injected mouse identified by the different symbols. Tumor sizes were measured every 2-3 days during the observation time. X-axis scale indicates the days of tumor monitoring, as well as the timing of tumor implantation, immunization and bleeding. B. Kaplan-Meier survival curve.

A good association between the levels of antigen-specific CD8^+^ T cell immune responses and antitumor effects was found in the group of mice injected with Nef^mut^/E7- and Nef^mut^/E6-expressing vectors. In fact, the mean of SFUs/10^5^ PBMCs measured in mice developing tumor was 115±61, whereas in cured mice increased to 187±92.

Taken together, these data demonstrated that the Nef^mut^/E6 plus Nef^mut^/E7 combined vaccine led to a tumor growth control resulting much more potent than that induced by the combined injection of DNA vectors expressing HPV16-E6 and -E7.

### Resistance of cured mice against tumor re-challenging

Next, we were interested in evaluating the persistence of the antitumor state in mice cured by the immunization with DNA vectors expressing Nef^mut^/E6 and Nef^mut^/E7. Six mice which were cured by the immunization were re-challenged by implanting 2×10^5^ TC-1 cells in the opposite flank respect to the previous cell implantation. As control, age-matched, non-immunized mice were used. We observed that cured mice remained tumor-free over the 135 days of monitoring, whereas the age-matched naïve control mice developed a palpable tumor after 12 days from TC-1 cell implantation, and had to be sacrificed at day 20 (Fig. 5A). The six tumor-free animals were sacrificed at the end of follow-up, and their splenocytes tested by IFN-γ Elispot assay to assess the presence of E6- and E7-specific CD8^+^ T cells. Detectable levels of both E6- and E7-specific CD8^+^ T cell immune responses were observed in all mice, in the presence of higher E7-specific immune responses (Fig. 5B). All re-challenged mice were confirmed tumor free and with normal-looking organs (i.e., lungs, heart, liver, spleen, kidneys, stomach, intestine) by observation at necroscopy.

**Figure 5.**
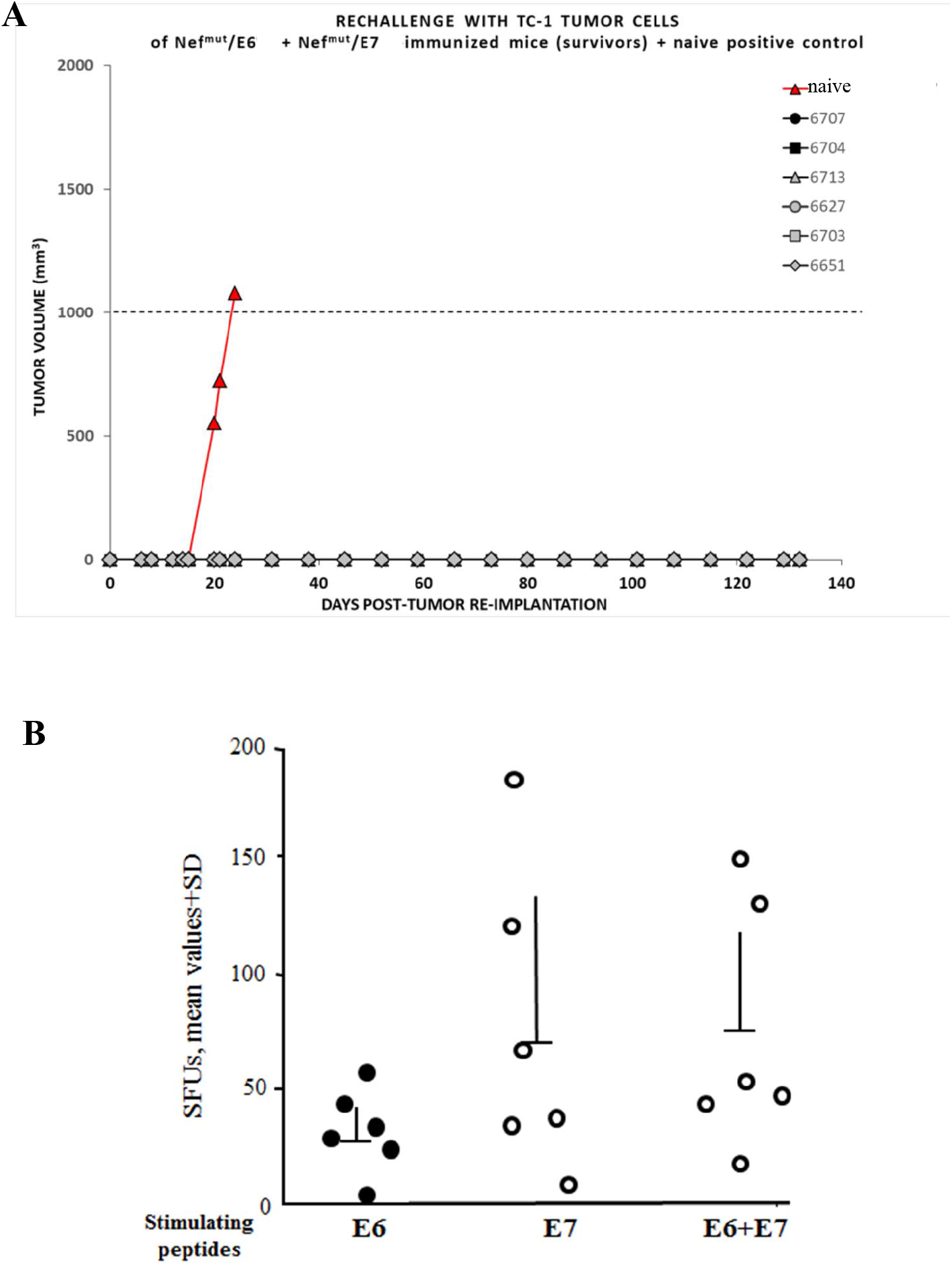
Antitumor therapeutic effect after TC-1 cell re-implantation. A. Tumor growth curves. Tumor-free C57 Bl/6 mice were re-challenged with 2×10^5^ TC-1 cells, and the growth of tumor mass was followed over the time. As control, age-matched naïve mice were inoculated with the same number of cells. Tumor sizes were measured every 2-3 days during the observation time. On the top left, the identification of each inoculated mice is shown. B. CD8^+^ T cell response in mice challenged, immunized, and re-challenged with TC-1 cells. Tumor-free C57 Bl/6 mice previously implanted with syngeneic TC-1 tumor cells and immunized with both Nef^mut^/E6- and Nef^mut^/E7-expressing vectors were re-challenged with TC-1 tumor cells. Nineteen weeks after cell re-implantation, mice were sacrificed, and splenocytes tested for both E6- and E7-specific CD8^+^ T cell immunity by IFN-γ EliSpot assay. Either E6, E7, or a mix of E6 and E7 peptides were used in triplicate microwells. The number of IFN-γ spot-forming units (SFU)/well are shown as mean values of triplicates. Intragroup mean values + SD are reported.

These results strongly suggested that the antitumor state induced by engineered EVs was strong and durable enough to counteract the proliferation of re-challenging tumor cells.

## Discussion

Duration and robustness of the immune response are key features for any vaccine strategy. The Nef^mut_^based CTL vaccine platform has been proven to induce strong immunity towards a wide range of both tumor and viral antigens (15, 16). However, log-term efficacy against tumor cell challenge and re-challenge not evaluated yet. To fill this gap, we first tried to identify the most efficient immunogenic strategy pertaining the Nef^mut^ technology. We found that co-injection of DNA vectors expressing different antigens fused to Nef^mut^ resulted in an additive anti-HPV16 CD8^+^ T cell immune response with no interference in terms of the downstream immunogenicity. Consistently, increased percentages of antigen-specific polyfunctional CD8^+^ T lymphocytes compared to those induced by single vectors have been observed. These results can be considered of great relevance since they open the way towards the application of the Nef^mut_^based platform on multiple targets in new vaccine combination strategies.

In tumor-implanted mice, the expression of fusion to Nef^mut^ of HPV16-E6 and -E7 antigens led to a strong increase of the antigen-specific CD8^+^ T cell immune response compared to that elicited by each HPV16 product expressed alone. Consistently, the antitumor effect was far more striking in mice injected with both DNA vectors expressing the products of fusion. Results from the tumor re-challenge experiment demonstrated that the Nef^mut^/E6 plus Nef^mut^/E7 vaccine conferred a potent, long lasting (at least up to 38 weeks after the last immunization), and efficacious CD8^+^ T cell immunity against the tumors re-implanted 19 weeks after the last immunization. At this time, immune responses against both HPV-E6 and E7 proteins were still detectable, and all mice remained tumor-free.

Taken together, these data prove that the vaccine strategy based on endogenously engineered EVs is a promising approach against HPV-antigen expressing tumors, and that it may be worth further testing against this and additional pathologies.

We previously described an antitumor therapeutic effect in mice injected with a Nef^mut^/E7 DNA vector (15). However, the tumor monitoring was limited to 30 days after tumor implantation, and the tumorigenicity of TC-1 cells appeared significantly reduced compared to that observed in the here presented experiments. In fact, at day 30 after tumor implantation all mice injected with control vectors survived and the tumor mass did not exceed 0.6 cm^3^, whereas at this time point in the here presented experiment all mice had to be euthanized.

In the classic mechanism of action underlying i.m. DNA vaccination, antigens expressed by DNA recipient muscle cells can be secreted, thereby essentially generating a humoral adaptive immune response (27). Muscle cells are not professional antigen presenting cells (APCs), and do not express co-stimulatory molecules. Hence, the CD8^+^ T immune response relies on the capture and expression of the DNA molecules by professional and semi-professional APCs (e.g., DCs, endothelial cells) embedded in the muscle tissue. However, naked DNA does not spread from cell to cell *in vivo*, and APCs do not efficiently take up exogenous DNA to activate satisfactory immune responses. For these reasons, the delivery of DNA vaccines in DCs is most effectively achieved through subcutaneous and intradermal injections, possibly associated with gold particles and gene gun. Differently from all currently developed DNA vaccines, in the Nef^mut_^based platform the expressed antigen is incorporated into the EVs, which are spontaneously released by the muscle cells, and are expected to freely circulate into the body. The Nef^mut_^based vaccine platform combines the remarkable benefits of efficient cross-presentation of EV-associated antigens and the consequent specific induction of a potent CD8^+^ T immune response, with several advantages typical of DNA vaccines, including: i) simple and flexible design, so that a wide range of antigens and immunomodulatory molecules can be expressed; ii) unrestricted MHC Class I immune response; iii) no unsafe infectious agents involved in the preparation of immunogens and, consequently, no adverse clinical effects or toxicity are expected to occur; iv) great heat stability and ease of storage and transport without need for a cold chain, and v) cost effectiveness. Immunogens can be developed quickly and easily once the antigen has been identified. The production can be very rapid, reproducible and perfectly suitable for large-scale production and administration.

Patient-derived EVs were employed as a novel cancer immunotherapy in several clinical trials, but this strategy lacked sufficient efficacy (28–30). Other lines of research have focused on modifying the content and function of EVs in various ways, toward the end-goal of specialized therapeutic EVs. Exosomes engineered to upload cargoes represent the last frontier in terms of nanoparticle-based technology (31), and a number of companies have been established on the basis of patented technologies to engineer EVs *in vitro*. Ideally, EVs could be designed for a desired immunostimulatory function and loaded with antigens or cell-targeting proteins *in vitro* to produce a potent, antigen-specific immune response *in vivo*, e.g. addressed to a functional tumor therapy. Despite high expectations, however, clinical trials have not yet confirmed therapeutic applicability of *in vitro* engineered EVs (32–34). Considering the underlying mechanism of action, the Nef^mut^ vaccine approach can overcome limitations pertaining DNA-based vaccines, as well as *ex vivo/in vitro* engineered EVs.

